# Calibration-free 3D reconstruction of firefly trajectories from 360-degree cameras

**DOI:** 10.1101/2021.04.07.438867

**Authors:** Raphaël Sarfati, Orit Peleg

## Abstract

Over the past few decades, progress in animal tracking techniques, from large migrating mammals to swarming insects, has facilitated significant advances in ecology, behavioural biology, and conservation science. Recently, we developed a technique to record and track flashing fireflies in their natural habitat using pairs of 360-degree cameras. The method, which has the potential to help identify and monitor firefly populations worldwide, was successfully implemented in various natural swarms. However, camera calibration remained tedious and time-consuming. Here, we propose and implement an algorithm that calibrates the cameras directly from the data, requiring minimal user input. We explain the principles of the calibration-free algorithm, and demonstrate the ease and efficiency of its implementation. This method is relatively inexpensive, versatile, and well-suited for automatic processing and the collection of a large dataset of firefly trajectories across species and populations. This calibration-free method paves the way to citizen science efforts for monitoring and conservation of firefly populations.

## Introduction

The considerable expansion of continuous, high-resolution, non-invasive animal tracking capabilities over the past few years is making a profound impact in ecology and animal behaviour science. Facilitated by new hardware and software technologies, tracking individual animals of various sizes, from insects to mammals, and across various length-scales, is allowing new insights into species distribution, migration patterns, response to changes of the environment, resource managements, social and collective behaviour, to name a few (*Kays et al., 2015*).

At the transcontinental scale, GPS devices, becoming smaller and less inexpensive (*Foley and Sillero-Zubiri, 2020*), have been placed on thousands of birds as well as terrestrial and aquatic mammals (*Wikelski, 2013*), providing a clear visualization of migration routes. In return, these results illuminate how climate change and a degrading environment influence migration timing and species resilience (*Middleton et al., 2013, Middleton et al., 2018*). They have also contributed to conservation efforts by pinpointing obstacles in migration corridors, such as fences and roads, so that animal crossing solutions could be implemented (*Middleton et al., 2020, Xu et al.,*). In parallel, computational techniques have been developed to manage and analyze these large datasets (*Wikelski, 2013, Dell et al., 2014*), and public data repositories, such as Movebank.org, have emerged for easy access by the scientific community.

At the habitat scale, three-dimensional (3D) trajectories of birds (*de Margerie et al., 2015*), fish (*Qian and Chen, 2017*), or insects (*Sinhuber et al., 2019*) obtained from stereoscopic video recordings have provided significant new understanding into how these animals behave collectively. Notably, they have revealed the emergence of group-level properties with significant biological and ecological consequences, for example for navigation (*Berdahl et al., 2013*), predator avoidance (*Storms et al., 2019*), and social interactions (*Butail et al., 2012, Ling et al., 2019*).

In parallel, 3D tracking allows to measure population densities, and hence to evaluate the resilience of potentially endangered species, such as fireflies (*Lewis et al., 2020, IUCN, 2021*). Recently, we developed a new experimental technique for tracking swarming fireflies using pairs of 360-degree cameras (*Sarfati et al., 2020*). The method is inexpensive, versatile, and easier to set up than traditional multi-camera systems. It is well-suited for imaging in complex natural habitats, such as dense forests, because 360-degree cameras record everything around them, so they can be placed directly within a swarm, rather than on the outskirt. It could also be extended to other animal collectives. The main limitation for a widespread use of this technique was the need for a tedious calibration operation, which was time-consuming and required advanced technical skills. In this paper, we present an improved algorithm which performs camera calibration from the data itself, so that virtually no user input is necessary. We present the principles of automatic calibration, their results on recordings of two firefly species in their natural habitat, and discuss possible applications. In particular, the new software could make large-scale citizen science efforts widely realistic. The code used for this paper is written in Matlab and is available at www.github.com/peleg-lab/stereo360_calibrationfree.

## Materials and Methods

### Field experiments

To film fireflies in their natural habitat during mating season, we used pairs of GoPro Fusion 360-degree cameras ($300 each) recording at 30 or 60 frames-per-second (fps). The settings were adjusted to record at a high ISO value of 1600 (the maximum value of 6400 appeared too noisy/grainy). Two cameras were positioned side-by-side in a cleared and flat area with high firefly activity, facing the same direction (Fig. 1a). The distance between them was set to either 0.9m or 1.8m (3ft or 6ft) using a flexible ruler. This step is indeed necessary to convert virtual units to physical dimensions. We studied two different species at two different locations: *Photuris frontalis* in Congaree National Park (CNP), South Carolina, USA, recorded in May 2020; and *Photinus carolinus* in Great Smoky Mountains National Park (GSMNP), Tennessee, USA, recorded in June 2020. These two species have different flash patterns: for *P. frontalis*, a fast, 10-30ms flash repeated every 0.7s 10 to 20 times (*Moiseff and Copeland, 2000*); for *P. carolinus*, a 100-150ms flash repeated every 0.6s 3 to 8 times (*Copeland and Moiseff, 1994*). Automated calibration is expected to work with most common flash patterns (see Discussion).

**Fig. 1.**
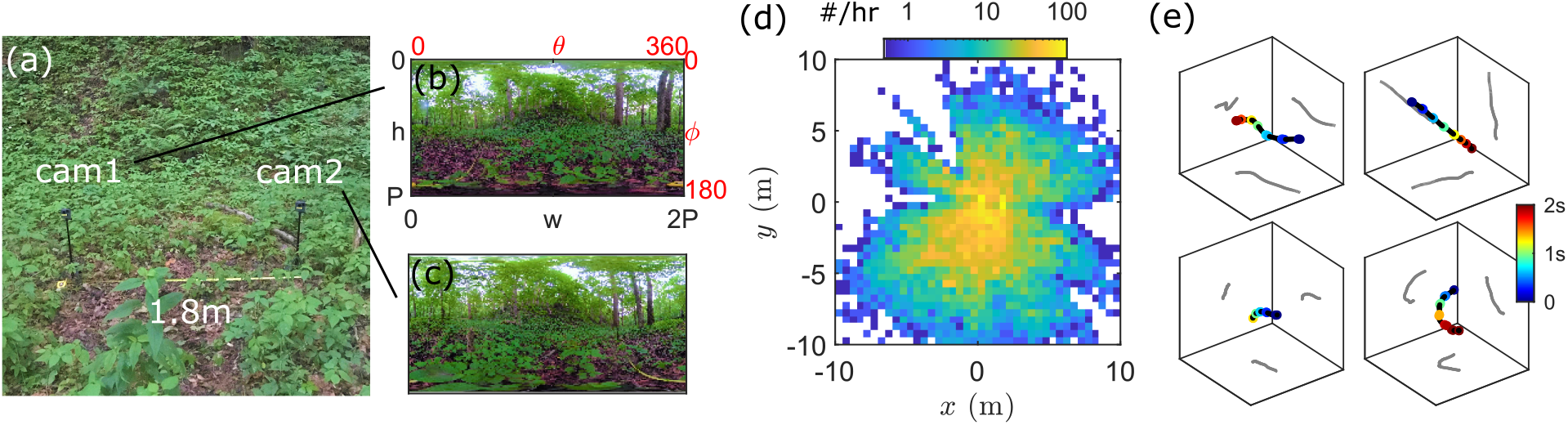
Experimental setup and 3D reconstruction results. (a) Pair of GoPro 360-degree cameras on their tripods (0.6m), facing the same direction and separated by a flexible ruler (1.8m). They were placed directly in the firefly natural habitat. Congaree NP, May 2020. (b,c) Complementary views from the left (b) and right (c) cameras. The photo-sphere is stereographically projected onto an equirectangular frame (dimensions 2*P* × *P* pixels^2^), which maps pixel locations (*w, h*) onto spherical angles (*θ, φ*). (d) 3D reconstruction of the flashing swarm, projected in the horizontal plane (‘viewed from above’). Colors indicate the number of flashes per hour (#/hr) in each spacial bin (0.5m)^2^. (e) Sample trajectories obtained after 3D reconstruction. Trajectories are obtained by extrapolation between successive streaks (colored dots).

### Principles of 3D reconstruction

The principles and implementation of 3D reconstruction from 360-degree cameras are exposed in details in Ref. (*Sarfati et al., 2020*). Briefly: 360-degree cameras record at 360°×180° around them, *i.e.* over the entire sphere. The photo-sphere is then rendered as an equirectangular frame of dimension 2*P* ×*P* pixels^2^. The planar coordinates (*w, h*) map onto the polar *θ* = 360° *·w /*2*P* and azimutal *ϕ* = 180° · *h/P* spherical angles (Fig. 1bc). Taking the location and orientation of Camera 1 as the coordinate system of the world, Camera 2 is translated by a vector *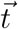* and rotated by a rotation matrix **R** relative to Camera 1. Let’s assume that a flash occurs at a position *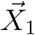* relative to Camera 1. Its projection in the field-of-view (FoV) of Camera 1 is (*θ*_1_*, ϕ*_1_), and in the field of Camera 2, (*θ*_2_*, ϕ*_2_). Based on these projections, 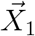 can be triangulated using the geometric relation:

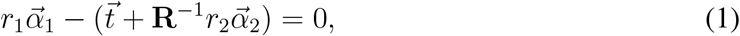

by solving for (*r*_1_*, r*_2_), with

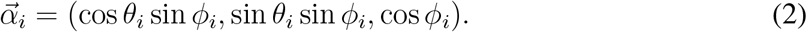

Hence, 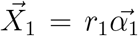. (Note that we set 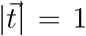, and conversion to real world units requires to rescale by the actual distance between the two cameras.)

Matching and triangulating flash positions requires to perform camera calibration, both in time and space. Specifically, one needs to measure the time delay, in number of frames Δ*k*, between the two cameras, as well as estimate the camera pose 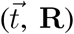. This calibration was previously done using an artificial flash signal for matching complementary frames, and the trajectory of a small light-emitting diode (LED) for spatial calibration. While relatively simple, these steps required additional experimental steps, and to play the movies, manually extract specific frames and identify the LED trajectory, which was time-consuming and involved significant human input. This was not convenient for automatic processing of many datasets coming from the large-scale deployment of the stereoscopic setup.

We later found that manual calibration could be avoided by relying instead on the data, *i.e.* recorded firefly flashes, from which 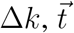, and **R** can all be inferred computationally. We present this updated method below.

Following automatic calibration and triangulation, flash occurrences can be placed in three-dimensional space, providing in particular density estimations (Fig. 1d). From 3D localization, full trajectories, consisting of several flashes, can even be inferred (Fig. 1e) (*Sarfati et al., 2020*).

## Automatic calibration from firefly data

### Temporal calibration

Automated frame synchronization relies on the cross-correlation of the time series of the number of detected flashes in each camera, *N*_1_(*k*_1_)*, N*_2_(*k*_2_), with *k_i_* the camera time expressed in frame number. Assuming first that these two traces are identical but shifted by Δ*k* = *k*_2_ − *k*_1_ frames, the cross-correlation will return the true value for Δ*k* (Fig. 2ab). In practice, however, the time series will be slightly dissimilar as they originate from different FoVs, where visual occlusion and limited light sensitivity impact what each camera records. With enough data, of the order of 10^5^ frames in a movie, this is not an obstacle. The most problematic source of noise comes from occasional background light pollution. Typically, this will occur at the beginning of the recordings, if the night is not yet obscure enough or if a flashlight is used to set up the cameras. (The latter might happen at the end as well.) Additionally, there are sometimes persistent objects in the FoV background, *e.g.* Moon, backyard light, passer-by with a flashlight, etc.

**Fig. 2.**
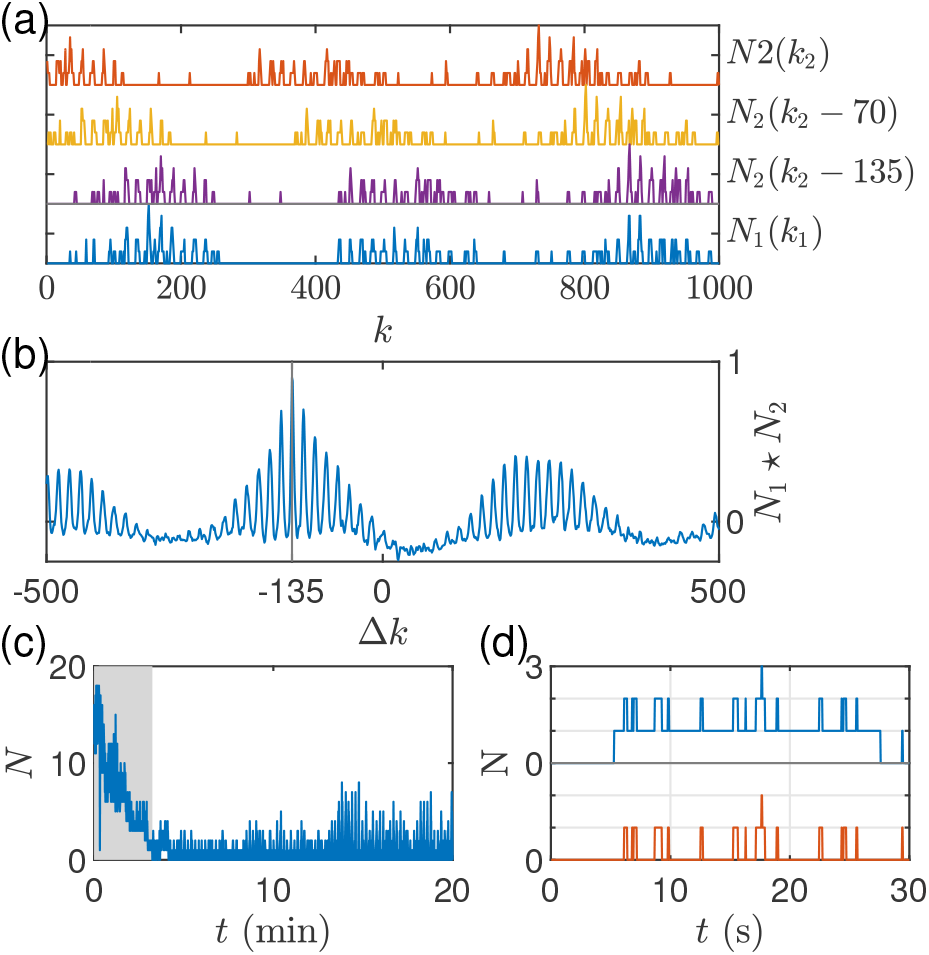
Temporal calibration. (a) *N*_1_(*k*_1_) and *N*_2_(*k*_2_) are short intervals of the time series of the number of flashes in camera 1 and camera 2, respectively; *k*_1_*, k*_2_ are the cameras’ internal times, in frame number. Also shown are successive shifts of *N*_2_ to align it with *N*_1_. (b) Corresponding cross-correlation *N*_1_ *\* N*_2_ spectrum, showing a maximum at Δ*k* = −135, which is when the two traces are best aligned in (a). (c) Example of large noise at the beginning of recording, due to sky illumination. The shaded area is automatically removed from the analysis. (d) Example of a persistent object in the field of view and its impact of the time series (top). Our algorithm removes these persistent objects (bottom).

To alleviate these issues, we start by pre-processing the original input, consisting of a set of (*w, h, k*) coordinates. The sets are truncated to remove the points before the first *N* = 0 and after the last *N* = 0 frames, which eliminates sky brightness and flashlight effects (Fig. 2c). Secondly, a simple tracking algorithm is applied to detect persistent objects lasting over a certain duration which would be impossible for a firefly to sustain, and therefore corresponds to light pollution (Fig. 2d). The threshold is set to 300 frames (5s or 10s, depending on frame rate), so as not to exclude glowing species, such as “blue ghost” fireflies (*Phausis reticulata*).

Finally, to make sure that the time delay estimation remains robust against possible residual noise, and estimate the reliability of the measurement, we apply a Random Sample Consensus-type algorithm (RANSAC) to the cross-correlation procedure: 100 random intervals are extracted from the cleaned traces, and cross-correlation is applied to every pair.

## Spatial calibration

Spatial calibration requires a set of broadly-distributed matched points so that 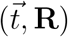 can be estimated according to the method described in Ref. (*Sarfati et al., 2020*). Originally, matched points were obtained from an identifiable trajectory, created manually with a small LED at the beginning of the recording (Fig. 3a). For the calibration-free algorithm, a set of matched points is collected simply by extracting all positions where *N*_1_(*k*_1_) = *N*_2_(*k*_2_ −Δ*k*) = 1 in both frames (Fig. 3). Most of the time, these flashes will indeed correspond to the same firefly, and will be a true matched point. Sometimes, however, they will come from two different sources. Consequently, the algorithm must be robust against outliers. For this reason, we employ a Maximum Likelihood Estimator for 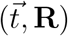. Taking as input a set of points, the algorithm calculates the values for 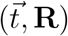 that maximize a likelihood function. As a global optimizer, outliers contribute little to the overall cost.

**Fig. 3.**
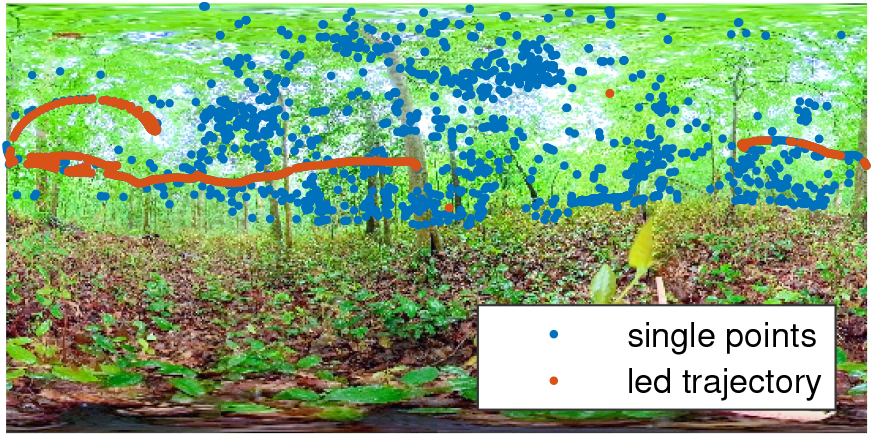
Spatial calibration. Originally, spatial calibration was performed using the set of points from the artificial trajectory of an LED (orange dots). For automatic calibration, the set of points instead consists of unique flashes detected in both FoVs (blue dots).

## Results

We produced 9 sets of stereoscopic recordings, each of about 90min, for both CNP and GRSMNP field experiments. We apply the automatic calibration algorithm on each of the 18 datasets, and compare the results to those obtained from manual calibration.

Regarding time delay estimations, we find an absolute agreement between manual measurements from flash signal and cross-correlation (Fig. 4a). In all but 1 dataset, the proper value was returned for 100% of the interval pairs. The remaining dataset had a success rate of 74% (Fig. 4b).

**Fig. 4.**
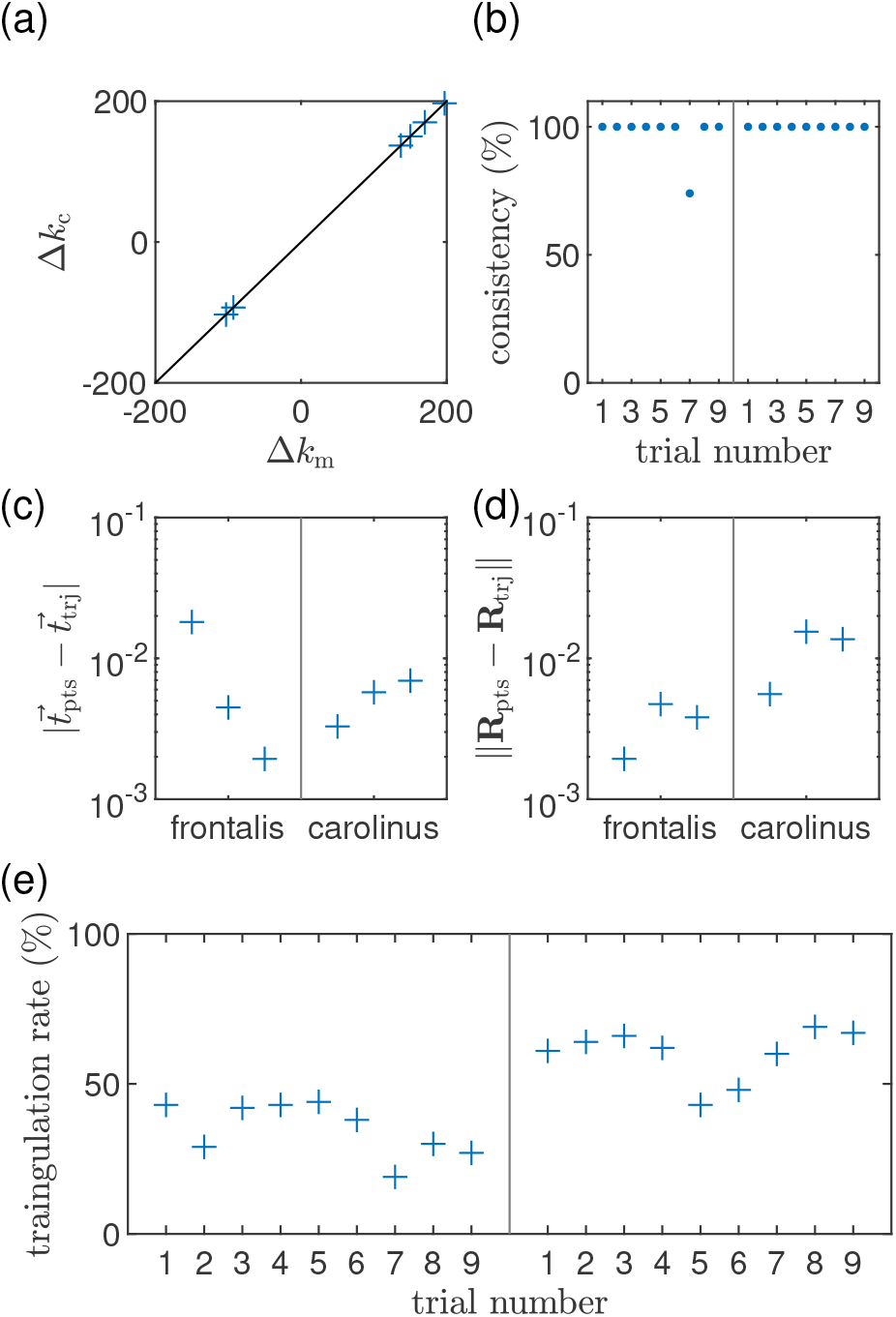
Calibration and triangulation results. (a) Comparison of frames delays obtained from manual identification of corresponding frames, Δ*k*_m_, and from computation of the cross-correlation spectrum, Δ*k*_c_. (b) Results from RANSAC time delay estimations. For each trial, 100 random intervals of the complementary time series are drawn, and cross-correlation is applied on each pair to estimate Δ*k*. All but one experimental set returns 100% consistency in using different time intervals. (c,d) Comparison of spatial calibration from unique points (pts), *i.e.* automatic calibration, and artificial LED trajectory (tra), *i.e.* manual calibration. The difference between translation vectors and rotation matrices obtained from both methods are of order 10^−2^ or less. The matrix 2-norm was used, and note that 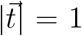 and det **R** = 1. (e) Triangulation rate from each trial, defined as the ratio between the number of 3D reconstructed flashes and the number of flashes in recorded separately in each camera. Due to visual occlusion and limited light sensitivity, triangulation rate is not expected to be significantly above 50%.

Regarding camera pose estimations, we compare the difference between translation vectors obtained from calibration using the LED trajectory (tra) and using single points (pts), and similarly with the rotation matrices. Specifically, we calculate 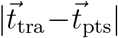 (Fig. 4c) and ∥**R**_tra_−**R**_pts_ ∥ using the matrix 2-norm (Fig. 4d). In all cases, the differences amount to a small discrepancy of order 10^−2^ or less.

Finally, a good metric of successful calibration is the ability to triangulate a large fraction of the points present in both cameras. Typically, due to visual occlusion from the surrounding vegetation and the cameras’ limited light sensitivity, many of the flashes captured in one camera may not be captured in the other, so that even a 50% rate should be considered successful reconstruction. If the calibration has failed, due to either an incorrect delay estimation or pose estimation, the triangulation rate is typically of order 1% or less. In Fig. 4d we report the triangulation rate for all datasets. These all lie between 30% and 60%, which is very satisfactory. When calibration from the LED trajectory is used in comparison, automated calibration performs identically (within 2%).

## Discussion

The imaging technique and associated algorithm presented here enable the 3D reconstruction of thousands of firefly positions and trajectories within a 10m-sphere around the cameras. The experimental procedure only requires to identify a good site, carefully position the cameras on the ground, and measure the distance between them. From the locations extracted from the movies (using for instance a thresholding method), our code then fully reconstructs the swarm *without any user input*. It also provides simply metrics to evaluate whether the reconstruction has been successful. This processing pipeline is well-suited for a fully-automated integration of data collected by multiple users, permitting the collection and analysis of a broad dataset.

For the calibration algorithm to be effective, however, certain conditions are necessary. In order to measure camera time delays from time series cross-correlation, a substantial number of flashes need to be recorded (minimum ∼ 10^3^). On the other hand, there needs to be enough unique flashes to obtain a large set for spatial calibration, so that extremely high firefly densities could produce inaccurate results. Given the reality of firefly flashing displays, this would however be unlikely to occur. Besides these two conditions, any flashing pattern that can be accurately captured by the cameras at a reasonable frame rate (30fps to 60fps) is expected to be successfully reconstructed.

Automated calibration opens up promising prospects for citizen science projects and large-scale monitoring efforts. Fireflies are much beloved insects (*Lewis et al.,*), from children and adults alike, and provide engaging material for science education (*Faust, 2004*). They are also rare and fragile beetles, displaying typically for a few days to a few weeks, and only 2-3hr hours at a time. Therefore, studying fireflies is a difficult enterprise for scientists. It requires to be at a specific place, at night, within a narrow season of the year, and most fireflies display during the same period. Therefore, studying different species, and different populations within each species, often requires many years of committed work, and results in limited datasets.

On the other hand, any careful observer, trained scientist or not, can identify firefly display, and report observations. When synchronous fireflies were first debated in the early 1900s, many people started writing from all over the United States about their own observations, in very localized areas (*Morse, 1916, Allard, 1916, Hudson, 1918*). A few hundred meters further, they may never have known about them. That is why taping into citizen science to document firefly display can become a powerful tool. Every curious citizen scientist, with about $600 worth of equipment (potentially sponsored or borrowed) could very easily contribute to a large database of firefly displays. From a collection of flash patterns and trajectories, significant further understanding could be made about firefly communication, evolution of flash signals, etc. Most importantly, 3D reconstruction would allow for a rigorous estimation of firefly density, hence permitting population monitoring. At a time when worldwide insect populations are at risk of collapse (*Wagner et al., 2021*), and fireflies may be amongst the most fragile (*Lewis et al., 2020*), a large-scale monitoring program could positively impact conservation efforts of a staple of the summertime experience of nature.

## Acknowledgments

We are very grateful to the National Park Service, Congaree National Park, and Great Smoky Mountains National Park for facilitating our research, and Becky Nichols, David Shelley, and Paul Super in particular. We are also very thankful to Lynn Faust, Julie Hayes, and Paul Shaw for encouraging conversations and experimental assistance.

## Authors contributions

RS and OP designed and performed research and wrote the paper. RS designed and implemented the code and analyzed data. All authors contributed critically to the drafts and gave final approval for publication.

